# Electron Paramagnetic Resonance Investigation of Nitrite Binding in Myoglobin

**DOI:** 10.1101/252775

**Authors:** Matthew Bawn, Fraser MacMillan

**Affiliations:** From the Henry Wellcome Unit for Biological EPR, School of Chemistry, University of East Anglia, Norwich, Norfolk, NR4 7TJ, UK

**Keywords:** hemoglobin myoglobin, spectroscopy, protein chemistry, small molecules, nitrosylation

## Abstract

It has been proposed that myoglobin (Mb) may act as a nitrite reductase under hypoxic conditions. Any mechanism describing such activity should take into account the binding geometry of the ligand to the heme. Crystal structures of horse-heart Mb and human hemoglobin-nitrite complexes suggest that the anion adopts an uncommon *O*-nitrito binding mode. Electron Paramagnetic Resonance (EPR) spectroscopy was employed to investigate the nature of nitrite binding to Mb at pH values ranging from 6.5 to 10.8. Results suggest that for ferric Mb at low pH, nitrite binds in the *O*-bound nitrito mode resulting in a low-spin (LS) iron center. Further a high-spin (HS) iron center is observed at high pH in Mb-Nitrite with spectral values different to that of purely HS-Mb that is proposed to be due to an *N-*bound nitrite. The yields of these two species were found to be influenced by pH.

**Background:** Myoglobin has been theorized to have a role as a nitrite reductase.

**Results:** *O*-bound nitrite produces a low-spin ferric heme complex, whilst at high pH a high-spin species is found proposed to be the *N*-bound form.

**Conclusion:** Nitrite may bind to heme in myoglobin via N-nitro or O-nitrito mode.

**Significance:** The mechanism of any nitrite reduction will depend on its binding to the heme cofactor.

## Introduction

It has been established that many mammalian heme proteins react with various nitrogen oxide species and that these reactions are biologically relevant to mammalian physiology. Recent studies have indicated a role for nitrite as an endocrine storage pool of NO^•^ that can be bio-activated along the physiological oxygen gradient in order to mediate a number of responses, including hypoxic vasodilatation, (1) cyto-protection following cardiac ischemia/reperfusion and NO^•^-dependant and independent signalling. It is generally believed that nitrite reduction by heme-containing nitrite reductase (NiR) enzymes begins by the ligation of the nitrite anion to the heme iron centre within these proteins. As such a determination of the binding mode adopted by the nitrite anion within the heme protein is an important step in the elucidation of the mechanism of NiR activity. It has been widely accepted for some time that the primary source of nitric oxide within biological tissue is the metabolism of L-arginine by NO^•^ synthase (NOS) (2). There is, however, increasing evidence for alternative NOS-independent mechanisms for NO^•^ synthesis. It is also believed that these mechanisms may function in situations where NOS-derived NO^•^ production is impaired (1,3). Inorganic NO_2_ is an endogenous anion produced by the oxidation of NO^•^ under aerobic conditions (4). Conversely, the opposite redox reactions yield NO^•^ in acidic conditions from the reduction of NO_2_. NO^•^ is thought to be generated via deoxy hemoglobin during blood circulation and to have vaso-dilatory properties.

This alternative mechanism for NO^•^ production may be of particular importance under ischemic conditions since the NOS mechanism is oxygen dependent. As a consequence of the renewed interest in nitrite ligation to Mb, Copeland and Richter-Addo (5) have investigated the crystal structure of horse heart Mb to analyze how nitrite interacts with the heme center. Although several binding modes are known for nitrite to metals (6), the coordination to hemes (7-9) has nearly always exhibited the common *N-*nitro mode (Fig. 1). The structures of synthetic iron-porphyrin compounds with both ferric and ferrous centers (10,11) have also revealed the *N-*nitro mode. In order to determine whether the binding mode of the nitrite anion was directed by the specific amino acid residues within the heme binding pocket Richter-Addo and colleagues performed crystallographic studies on a series of Mb variants (12). They observed that when hydrogen bonding residues were present the *O-*nitrito mode was preferentially adopted by the ligand and when these residues were removed the *N-*nitro mode was more favored. Continuous Wave (cw)-Electron Paramagnetic Resonance (EPR) spectroscopy has been shown to be an excellent spectroscopic tool for the investigation of the heme-ligand structures in both low spin (LS) and high spin (HS) ferric heme systems (13). These spin-states produce predominantly either rhombic or axial EPR spectra, respectively, the g-values of which provide an indication of the local symmetry within the heme moiety.

**Figure 1.**
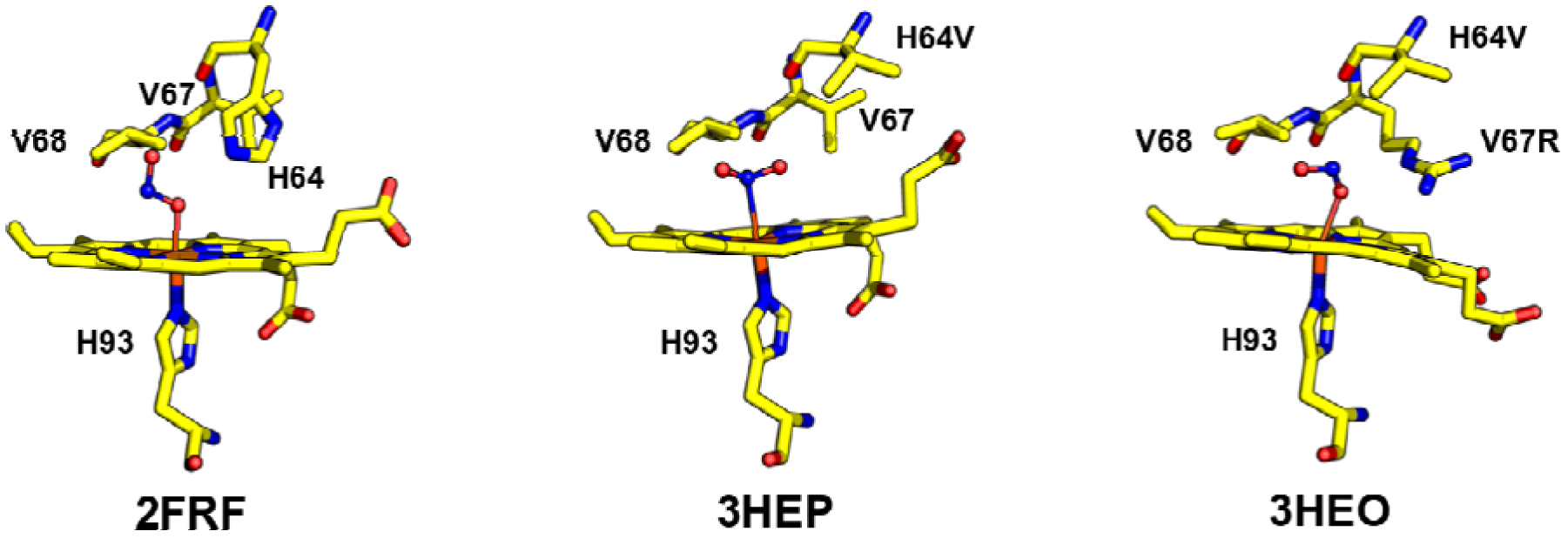
Heme center from crystal structures of Mb with ligated nitrite anion. (2FRF; native Mb, 3HEP; Mb with H64V mutation and 3HEO; Mb with H64V and V67R mutation). Structures demonstrate *O-nitrito* (2FRF and 3HEO) and *N-nitro* (3HEP) binding modes.

To determine affect of hydrogen bonding to the binding mode adopted in ferric Mb unconstrained from the restrictions imposed by crystallisation, EPR spectroscopy was applied to a series of Mb-NO_2_ complexes over a range of pH values (from 6.5 to 10.8).

## EXPERIMENTAL PROCEDURES

### EPR Sample preparation

Mb samples were made using horse heart Mb, all reagents were purchased from Sigma-Aldrich and used without further purification. 50 mM MES, HEPES, TAPS CHES and CAPS buffers were used to obtain samples of pH 5.5, 7.5, 8.5, 9.5 and 10.8 respectively. Phosphate buffers were avoided due to the known pH artifacts associated with freezing (14). Nitrite was added in the form of sodium nitrite to 240-fold excess. Na^14^NO_2_ was of at least 97% purity with sulfite and chloride anion concentrations of less than 0.01% and 0.05% respectively. For isotope labeling experiments Na^15^NO_2_ contained 98% Na^15^ and was 95% complete product isotope labeled nitrite. Samples were contained in suprasil quartz EPR tubes and stored at 77 K when not in use.

*cw-EPR spectroscopy* was performed using a *Bruker* eleXsys E500 X-band (nominally 9.4 GHz) spectrometer fitted fitted with a ER 4122 SHQ cavity and an *Oxford Instruments* (Oxford Instruments, Oxfordshire, UK) ESR 900 continuous flow helium cryostat. Spectra were recorded at 10 K using a microwave (mw) power of 0.1 mW a 100 KHz modulation frequency and a modulation amplitude of 5 G. Magnetic field values were calibrated against a DPPH standard. The mw frequency was measured using a Marconi Instruments 2440 mw counter.

*Pulsed EPR spectroscopy* was performed on a *Bruker* eleXsys E680 spectrometer fitted with a *Bruker* MD5-W1 resonator and equipped with an *Oxford* helium CF 935 flow cryostat. For LS samples spectra were measured at 10K and for HS samples the temperature was lowered to 4 K.

### Field-Swept Echo (FSE) spectra

Samples were initially characterized by measurement of the FSE spectrum (the pulsed-EPR equivalent of a cw-EPR experiment) prior to more advanced pulsed experiments. A*π/2* – *τ* – *π* – *τ* pulse sequence is applied as the magnetic field is swept in order to obtain an EPR spectrum.

### Three-pulse Electron Spin Echo Envelope Modulation (ESEEM)

The three-pulse ESEEM pulse sequence of *π/2* – *τ* - *π/2* – *T* - *π/2* – *τ* was applied in conjunction with ^15^N labeled nitrite to determine the hyperfine coupling from nitrite nitrogen in the LS complex.

### Matched HYSCORE

matched-HYSCORE was applied in order to optimize nitrogen hyperfine couplings, it (15) incorporates matched pulses (16) into the standard HYSCORE sequence. A matched also known as a high-turning angle (HTA) pulse increases the efficiency of the forbidden transfers within the experiment in order to achieve larger signal intensities in both ESEEM (Electron Spin-Echo Envelope Modulation) and HYSCORE experiments. A *π/2* – *τ* – *HTA* – *T*_*1*_ – π *T*_*1*_ – *HTA* – τ pulse sequence was used with HTA pulses of 128 ns. The application of matched pulses to the HYSCORE experiment has been shown to increase the signal-to-noise ratio for strongly coupled nitrogens significantly. The sequence has also facilitated the application of the HYSCORE technique to S > 1/2 systems (17).

### Hyperfine Sublevel Correlation (HYSCORE)

The standard HYSCORE pulse sequence *π/2* – *τ* - *π/2* – *T*_*1*_ – π – *T*_*2*_ - *π/2* – *τ* (140 ns τ-value, 16 ns π/2 and 32 ns π pulse) was used to examine the contribution to the Mb and Mb-NO_2_ HYSCORE spectrum from protons in the HS complex.

### Simulation of EPR spectra

Spectra are simulated using the *EasySpin* (18) toolbox for Matlab.

## RESULTS

### Increase in HS signal of Mb with addition of nitrite

The EPR spectra obtained for 2 mM Mb and Mb-NO_2_ at pH 10.8 are presented The spectrum of isolated Mb at pH 10.8 (Fig. 2A), is dominated by a LS rhombic signal associated with the Mb-hydroxide complex (19). A very small contribution from a HS signal, attributable to aquomet-Mb, is also observed. Typical HS (S = 5/2) heme EPR spectra are predominantly axial, having a low-field component at g_x,y_ ≈ 6 and components at g_z_ ≈ 2. The addition of nitrite to the same sample (Fig.2B) led to the observation of an additional LS rhombic EPR signal with g-values [g_ZZ_ ≈ 3.03, g_YY_ ≈ 1.53] (following the axis assignment for LS Heme complexes of Walker (13)) which are very comparable to those values observed previously for a Mb-nitrite complex (19). At this pH there is, however, still a contribution from Mb-hydroxide [g_ZZ_ ≈ 2.62, g_YY_ ≈ 2.18 g_XX_ π 1.86]‥A closer examination of the HS EPR signal after addition of nitrite reveals an increase in intensity of the low-field component and a change of g-values from [g_x,y_ ≈ 5.95, g_z_ ≈ 2.015] for the aquomet-Mb signal to [g_x,y_ ≈ 5.99, g_z_ ≈ 2.00] after the addition of nitrite.

**Figure 2.**
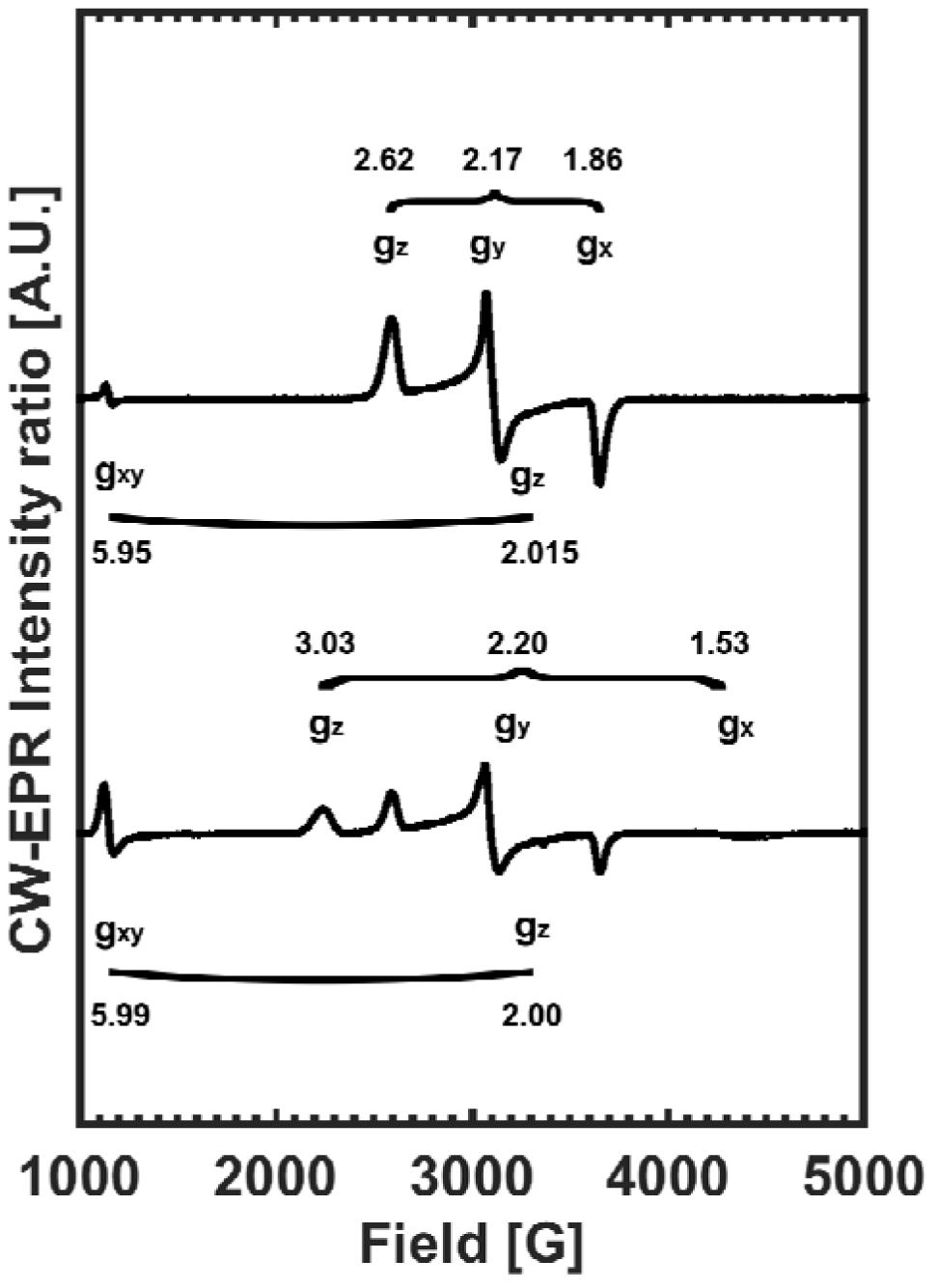
10 K EPR spectra of 2 mM Mb in CAPS buffer (pH 10.8) before and after the addition of 240-fold molar excess nitrite (top and bottom respectively). g-values of each species indicated.

### Quantification of species contribution from spectral simulation

samples of 250 each pH value were measured at 10 K. Samples were then thawed, had nitrite added and were remeasured (Fig.3A-E). The spectra were then simulated using g-values for each species as indicated above. The change in the contribution to the spectra with the addition of nitrite were then plotted as a function of pH (Fig. 3F-G). At pH 6.5 and 7.5 Mb is initially found in a purely HS state. Addition of nitrite greatly reduces the HS signal and yields a LS Mb-NO_2_ spectrum with g-values slightly different to those observed at pH 9.5 and 10.8 ([g_ZZ_ ≈ 3.0 g_YY_ ≈ 2.21 g_XX_ ≈ 1.56]). At pH 8.5 Mb is found in a predominantly HS state but with a LS. Nitrite addition also greatly reduced HS contribution and led to a LS Mb-NO_2_ spectrum with g-values as seen at pH 6.5 and 7.5. For pH 9.5 and 10.8 Mb was predominantly in a LS form. An increase in HS EPR signal was found after nitrite was added as well as LS Mb-NO_2_ and LS-Mb. At pH 10.8 the contribution to the spectrum from the HS signal increased approximately two-fold after nitrite addition.

**Figure 3.**
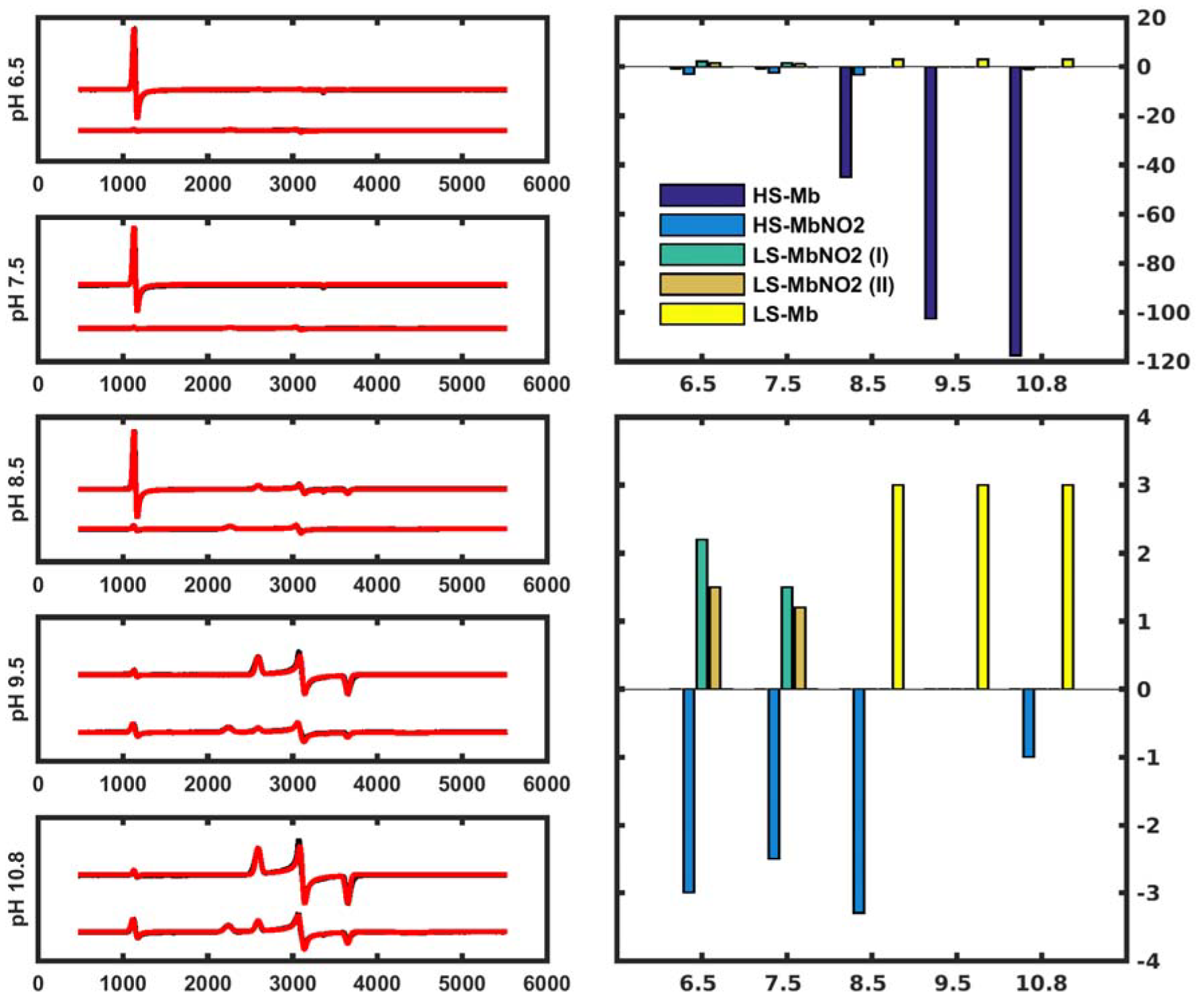
CW-EPR spectra (black) and simulations (red) measured at 10 K of 250 μM Mb before and after addition of 240-fold molar excess of nitrite (A - E) at pH 6.5, 7.5, 8.5, 9.5 and 10.8. Change in contribution from each species as determined from simulation (F and G). Plot G is as plot F but with the HS-Mb contribution removed to more clearly show the contributions from the other species.

### Three-pulse ESEEM of ^15^N nitrite

The 10 K spectra of 2 mM ^14^N and ^15^N Mb-NO_2_ pH 7.5 (Fig.4a) measured at cannonical positions for the LS-NO_2_ case, revealed spectral changes associated with an isotope effect only at the g_z_-position. Simulation of the time and frequency domains of this position (Fig.4B) yielded the following parameters; [-0.05 -0.05 4.03] MHz hyperfine coupling and [-0.2475 -0.25025 0.5] MHz quadrupole couplings for the nitrite ^14^N.

**Figure 4.**
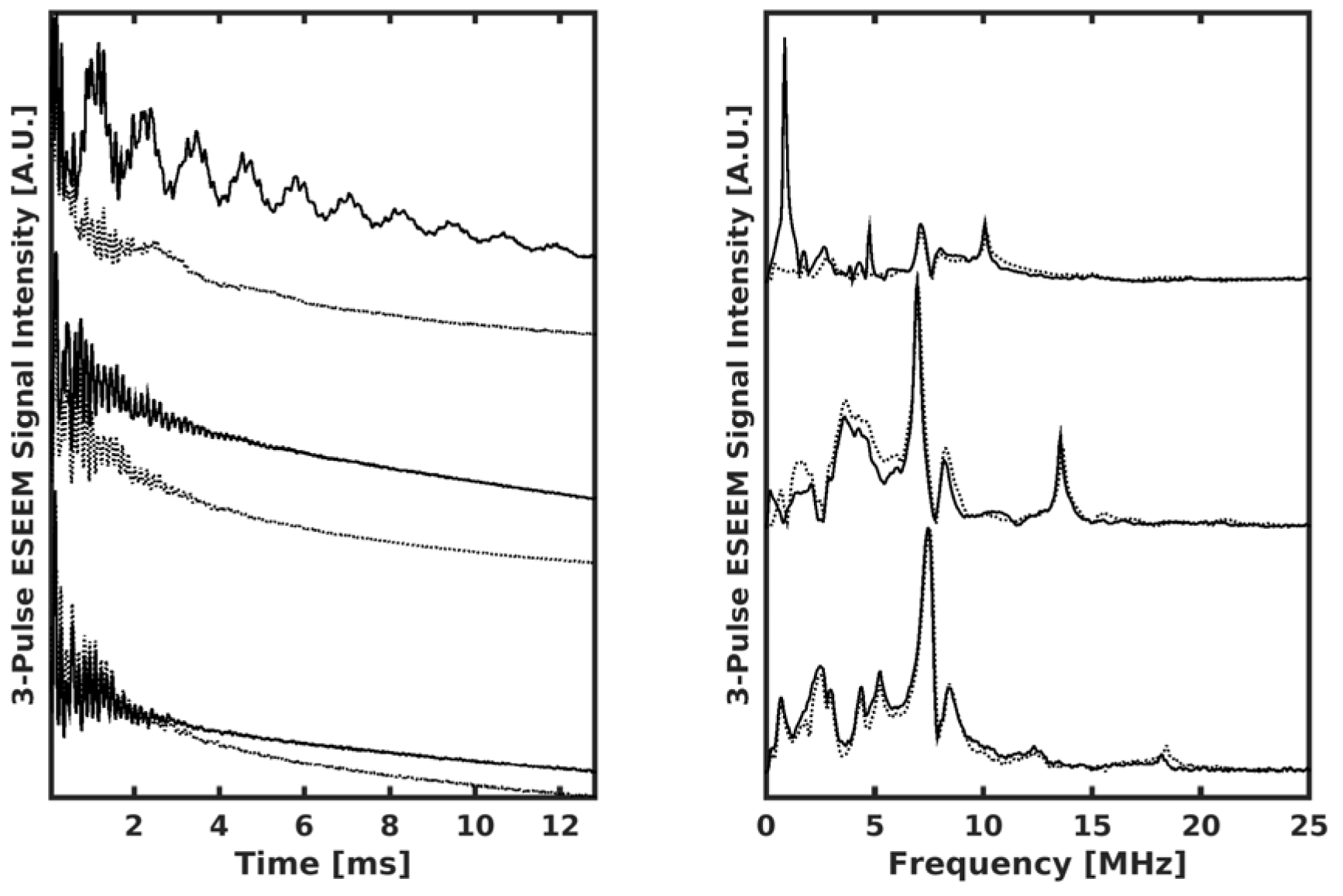
Time-domain and frequency-domain (A and B) three-Pulse ESEEM spectra of Mb- ^14^NO_2_ and Mb-^15^NO_2_ (Solid and dotted lines) measured at canonical positions. Spectra measured at 10 K.

### Matched-pulse HYSCORE

The matched-Pulse HYSCORE spectrum of LS Mb-^14^NO_2_ at pH 7.5 recorded at 10 K is shown (Fig 5). After Fourier Transformation of the two dimensional data set the figure consists of two quadrants, [–,+] and [+,+] which represent domains where strong(A_iso_ > 2v_14N_) and weak (A_iso_ < 2v_14N_) hyperfine couplings respectively are observed. In the [–,+] quadrant peaks which are associated with the double-quantum (DQ) transitions of the directly coordinating heme and histidine (His93) nitrogens are observed. A contribution from the ^14^N of nitrite is apparent in the [+,+] quadrant, whose magnitude is consistent with a nitrogen atom that is not directly coordinated to the Fe atom (20).

**Figure 5.**
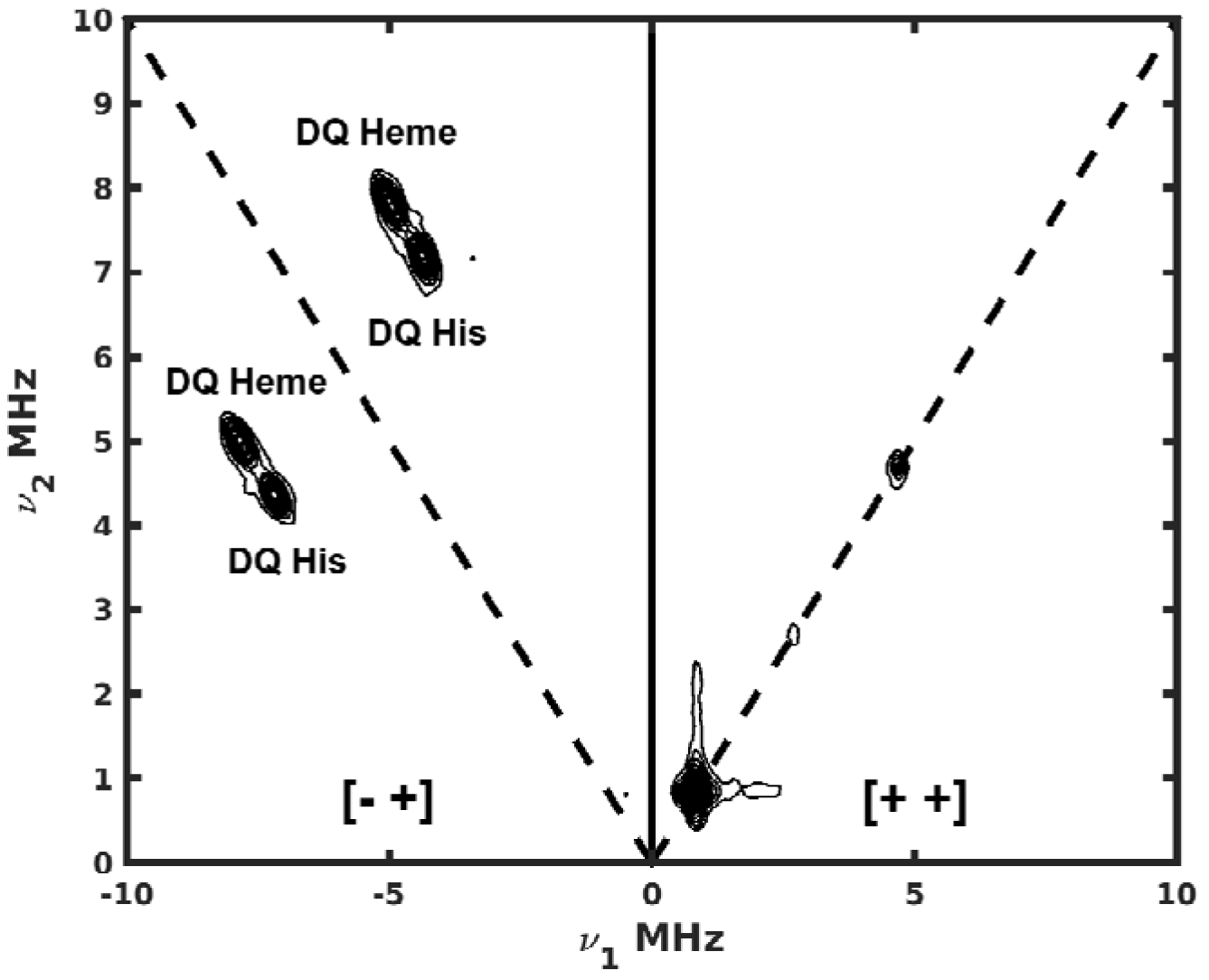
Matched-pulse HYSCORE of 2mM Mb-NO_2_ (pH 7.5) measured at 10 K. Contribution to spectrum from the nitrite nitrogen is circled with a dashed line in the [+, +] quadrant.

### FSE of HS complexes

The 4 K FSE spectra of HS Mb and Mb-nitrite at pH 7.5, 9.5 and 10.8 measured, a (τ-value of 160 ns and 32 ns π-pulse), are shown (Fig. 6A). For these spectra measured using pulsed-EPR the position of the low-field g-value appears at different positions for the samples examined. Hs Mb and Mb-nitrite at pH 7.5 have a peak at around g = 5.86 whilst the Mb-nitrite samples at pH 9.5 and 10.8 have this peak at a visibly lower g-value of 5.48. It may also be seen that the other spectral features are located at the same positions in each sample (e.g. the g = 2 peak) which rules out a field-offset between the spectra being responsible for the g-value difference.

**Figure 6.**
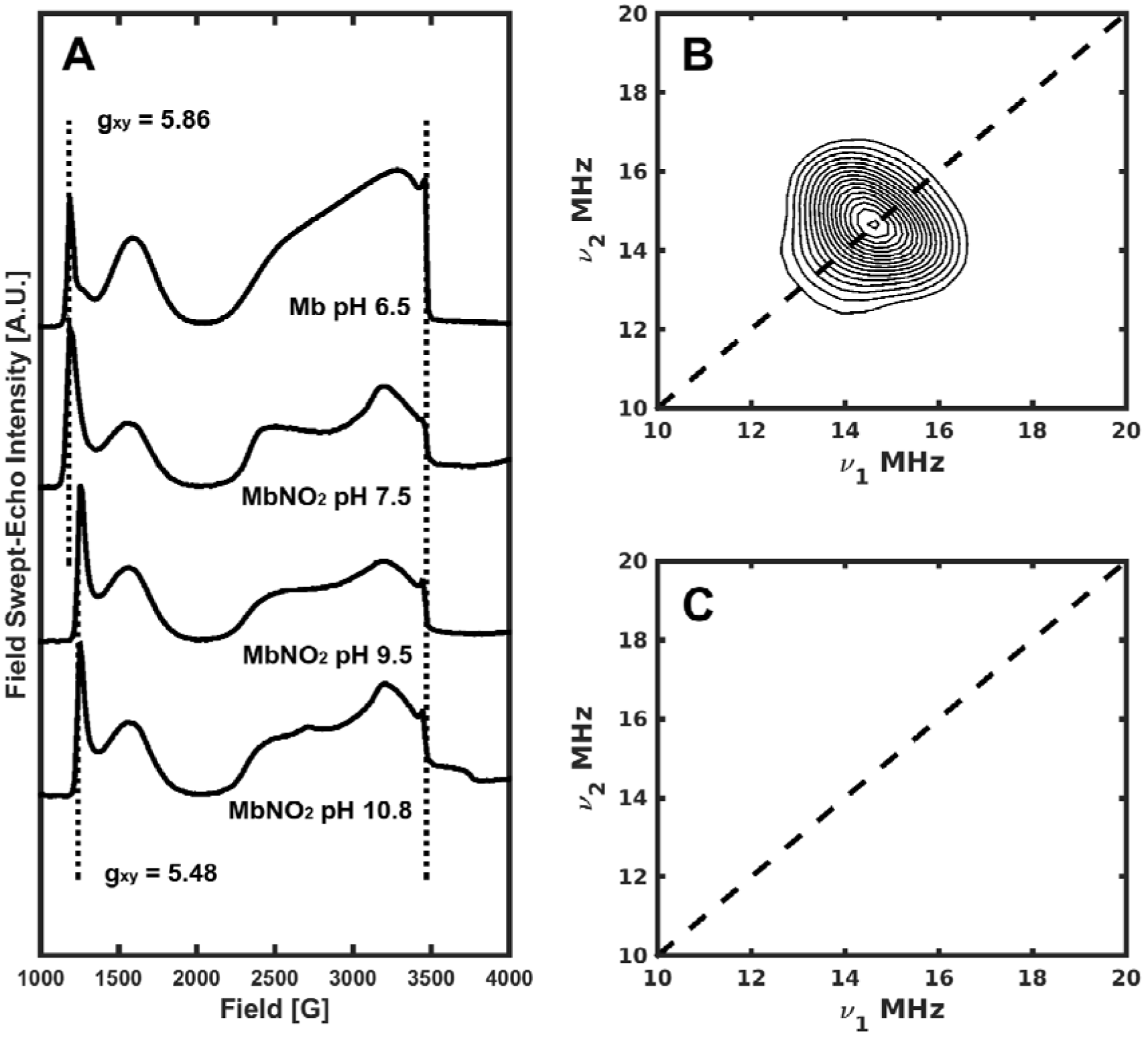
FSE spectra of a series of Mb and Mb-NO_2_ complexes as indicated (A). HYSCORE spectra of Mb at pH 6.5 (B) and Mb-NO_2_ pH 9.5 (B). Spectra measured at 4K. (B) shows the contribution of proton couplings absent in (C).

### Proton HYSCORE of HS Mb-NO_2_

The standard HYSCORE spectra of HS-Mb (pH 6.3) and the HS contribution from Mb-nitrite (pH 9.5) (Fig 6B and 6C) were measured at 4 K at the g_z_^eff^ position (~ 3440 G). In the HS-Mb spectrum the correlation associated with a weakly coupled proton is seen centered at the proton Larmor frequency (~ 14.65 MHz). this correlation has been previously reported and assigned to the protons of a coordinated water molecule with a hyperfine coupling at this position of ~ 5.9 MHz (20), this value also agreed with the seminal single-crystal ENDOR study of Scholes (21). The figure also shows the HYSCORE spectrum of Mb-nitrite at pH 9.5 measured with the same experimental parameters and conditions that favor the measurement of the HS signal. The spectrum was also measured at the g_z_^eff^ position. In this spectrum proton contributions are absent suggesting that the HS species does not have a coordinated water or even hydroxide molecule.

## DISCUSSION

### Increased HS-signal with nitrite addition

The observed increase in the HS contribution at high pH is unlikely to be due to a coordinated water complex since the pk_a_ of water coordinated to ferric Mb is known to be 8.9 (22). Above this pH the aquo-ligand is more likely to be deprotonated which results in a LS EPR signal. Interestingly a similar study of nitrite binding in hemoglobin (Hb) has revealed significant HS signals at high pH although the origin of these signals was not discussed (23). Richter-Addo *et al’s* findings suggested that the O-bound mode was facilitated by hydrogen bonding residues in the heme binding pocket, the most significant residue being His64. The pK_a_ of His64 is 4.4 (24), and is unlikely to be protonated above pH 8. One possible explanation for the increased HS signal, observable above pH 8.5,could be the presence of the *N*-nitro mode; hence the LS signal associated with the addition of nitrite would therefore be due to the *O*-nitrito form. In principle pulsed EPR spectroscopy allows the observation of weak hyperfine interactions that are masked in the broad cw-EPR spectrum to be resolved (25). Pulsed-hyperfine EPR spectroscopy in particular, can provide a detailed insight into the interactions of nearby magnetic nuclei with a paramagnetic centre.

### Examination of LS nitrite complex

Three-pulse Electron-Spin Echo Envelope Modulation (ESEEM) spectroscopy (26) was employed in conjunction with ^15^N isotope labeled nitrite to observe the existence of a hyperfine contribution from the nitrite nitrogen atom in the LS case. An accurate determination of the complete ^14^N/^15^N hyperfine interaction allows the distinction between the two possible modes of nitrite binding. Both time and frequency-domain spectra of Mb-nitrite (at pH 7.5) are depicted (Fig. 4A). In the time domain spectra (left) a clear effect of isotope substitution is observed when the field position chosen for the ESEEM experiment is aligned along the g_ZZ_ component of the LS heme signal. An apparent oscillation is now present which results, after Fourier Transformation, in an intense peak at ≈ 1.5 MHz (right, top trace). A spectral simulation of these spectra yielded the following parameters; [-0.05 -0.05 4.03] MHz hyperfine coupling and [-0.2475 -0.25025 0.5] MHz quadrupole couplings for the nitrite ^14^N. The magnitude of this ^14^N hyperfine coupling is rather small and not typical of a directly coordinated N-atom. A large hyperfine coupling constant is expected for a directly coordinated nitrogen atom. In order to resolve the observed nuclear hyperfine interactions further HYSCORE spectroscopy (27) was employed. A standard HYSCORE pulsed sequence employed along the g_ZZ_ orientation was poorly resolved, hence matched-pulse HYSCORE was employed. The observation of the nitrite nitrogen coupling peak being present in the [+,+] quadrant was indicative of a indirectly coordinated atom. This assignment supports the interpretation of the LS species being *O*-bound to nitrite.

### Investigation of HS signal

Pulsed-EPR was also applied to investigate the HS species formed after addition of nitrite. In order to obtain the FSE spectra of Mb (pH 6.5) and Mb-nitrite (at pH 7.5, 9.5 and 10.8) (Fig. 6A), a temperature of < 4 K was required. Contributions from any LS species were suppressed further by optimizing the microwave power used in the Hahn-echo sequence for these HS species. The two panels on the right (Fig 6B and 6C) also present the ^1^H-region of the respective HYSCORE spectra recorded at the g_ZZ_-position for Mb at pH 6.5 and Mb-nitrite at pH 9.5. The same pulse-lengths and distances were the same as used in the FSE spectra. The top two spectra (Fig. 6A) are recorded at low pH and reflect either a purely HS-Mb state (aquomet Mb) or this state in the presence of nitrite and where it is expected that a water molecule is coordinated to the iron center. HYSCORE spectra recorded at g_eff_ (g~2) reveal a correlation centered at the proton Larmor frequency. The bottom spectra are Mb samples in the presence of nitrite at higher pH (9.5 – 10.8) where the novel HS species is observed. The respective HYSCORE spectra (Fig. 5C) indicate that under these conditions no ^1^H correlations are observed, which would suggest that this is a new HS species formed by the addition of nitrite, as it does not contain a bound water molecule as found for HS-Mb. It is proposed that this species arises from an *N*-nitrito HS-Mb species, although the presence of a spin-equilibrium cannot be ruled out. In addition, as with the CW-EPR spectra there is also a measurable variation in the g-value of the lower field features of these two states from a g-value of 5.48 (at both pH 10.8 and 9.5) to a g-value of 5.86 at low pH. The difference is more marked in pulsed EPR however, due to tau-value enhancement by the filed-swept echo. These results clearly suggest that this new HS species observed at high pH is not due to coordinated water or hydroxide and is therefore likely to be due to coordinated N-bound nitrite. Indeed, in a recent publication density functional theory suggested both nitrite forms to be energetically favorable (28).

There is evidence that the dynamics of hydrogen-bonding may be related to enzymatic activity in at least one other heme protein (29) and hydrogen-bonding is responsible for differing oxygen binding energies in Mb and Hb from different species (30). It is therefore likely that any nitrite reductase activity of Mb may be facilitated by differential ligation of the ion to the heme center controlled *via* hydrogen bonding.

## Footnotes

* This work was financially supported by The Royal Society. FM is a Royal Society Wolfson Research Merit Award holder. EPSRC is also thanked for funding a PhD studentship.

3 The abbreviations used are: EPR, Electron Paramagnetic Resonance; Mb, Myoglobin; Hb, Hemoglobin; NiR, Nitrite Reductase; LS, Low-Spin; HS, High-Spin; CW, Continuous-Wave; ESEEM, Electron-Spin Echo Envelope Modulation; HYSCORE, Hyperfine Sublevel Correlation; DQ, Double-Quantum.

